# A hidden thermal mechanism in inhibitory ligand-gated chloride channels

**DOI:** 10.64898/2026.03.15.711968

**Authors:** Kohei Ohnishi, Yuichiro Fujiwara

## Abstract

Temperature profoundly shapes neural activity, with temperature-sensitive excitatory cation channels serving as key molecular components of thermosensation and neuronal excitability. Here we show that the glutamate-gated chloride channel AVR-14B, a Cys-loop receptor and principal molecular target of the antiparasitic drug ivermectin in the nematode *Brugia malayi*, exhibits temperature-dependent gating. Application of glutamate to AVR-14B evoked rapidly desensitizing transient currents; however, above approximately 24 °C, an additional non-desensitizing sustained current component emerged. Mechanistically, warming altered the gating behavior of the channel, thereby conferring intrinsic temperature sensitivity. Mutational and structural analyses revealed that ions permeate via lateral fenestrations distinct from the central axial pore, forming a noncanonical pathway for temperature-dependent gating. This temperature-dependent gating determines drug efficacy: ivermectin failed to activate AVR-14B below the thermal threshold at which the sustained current emerges. Finally, AVR-14B–null *C. elegans* showed enhanced heat tolerance, even though wild-type animals generally fail to thrive above 25 °C, confirming that this molecular mechanism governs organismal physiology. Similar temperature-dependent gating was observed in the human glycine receptor, indicating a conserved principle within this receptor class. Our findings identify inhibitory ligand-gated ion channels as intrinsic thermosensors and uncover a mechanism by which temperature can switch a single receptor between phasic and tonic inhibition, with implications for neural function and temperature-mediated therapeutics.

## Main

Temperature is a fundamental physical variable that profoundly influences neural activity and behavior across species. Understanding how neural circuits sense and integrate thermal fluctuations remains a central challenge in sensory physiology and neuropharmacology^1^. Thermo-sensitive TRP channels expressed in peripheral sensory neurons are well established as molecular thermosensors^2-4^. More recently, AMPA-type glutamate receptors—key mediators of excitatory synaptic transmission in sensory neurons—have also been shown to exhibit temperature sensitivity^5^, highlighting an emerging paradigm of temperature-dependent modulation among excitatory ion channels. Pentameric ligand-gated chloride channels—including glutamate-gated chloride channels (GluCls) in invertebrates and glycine receptors (GlyRs) in vertebrates—are evolutionarily conserved and critical for motor coordination, sensory processing, and reflex control^6-8^. GluCls are also the primary molecular targets of ivermectin, a landmark antiparasitic drug that has saved millions of lives and reshaped global health through sustained activation of these channels, inducing paralysis in nematodes^9-14^. Despite extensive structural and pharmacological characterization of these inhibitory channels, their potential role as intrinsic thermosensors—and the mechanisms that might support such function—has never been explored. This gap obscures our understanding of inhibitory physiology under varying thermal conditions and complicates the interpretation of temperature-dependent drug responses. GluCls and GlyRs both belong to the Cys-loop receptor family, whose members have recently been shown to harbor lateral fenestrations in addition to their central axis pores^15-18^. These lateral fenestrations are hypothesized to support alternative ion permeation pathways, but direct functional evidence has been lacking.

In this study, we demonstrate that the GluCl AVR-14B from the human-parasitic filarial worm *Brugia malayi* (*B. malayi*)^19-21^ exhibits intrinsic thermosensitivity with activation thresholds matching cultivation temperatures. By conducting structure–function analysis, assessing the temperature-dependence of pharmacological responses, and performing behavioral assays of thermal tolerance in nematodes, we reveal how temperature modulates gating and pharmacological responsiveness in Cys-loop receptors. Our findings establish inhibitory ion channels as previously unrecognized targets of thermal modulation, and highlight the functional importance of noncanonical permeation pathways, including lateral fenestrations. This work expands the conceptual framework of thermosensation in the nervous system and provides a novel foundation for therapeutic strategies.

## Results

### Thermosensitivity of *Bma-*AVR-14B channel

The *B. malayi* GluCl *Bma*-AVR-14B was expressed in *Xenopus laevis* oocytes and analyzed by two-electrode voltage-clamp recording. Currents evoked by the application of 1 mM L-glutamate were continuously recorded from a single oocyte at 15, 20, 25, 30, and 35 °C while the membrane potential was held at −80 mV in a Ca²⁺-free ND96-based solution (98 mM NaCl, 2 mM KCl, 2 mM MgCl₂, and 5 mM HEPES; pH 7.4). The recording temperatures were controlled by perfusing the oocyte with pre-warmed or pre-chilled solutions and monitored near the oocyte using a thermistor. At 15–20 °C, application of L-glutamate evoked a rapidly desensitizing transient inward current, consistent with prior descriptions of *Bma*-AVR-14B^22^ (Fig. 1a). Increasing temperature progressively enhanced the current amplitude, and above 25 °C we observed the emergence of an additional sustained component that continued to increase with further warming (Fig. 1a). Recordings from multiple oocytes showed that, when both current components were normalized to the amplitudes measured at 20 °C, the sustained current at 35 °C reached on average nearly ten times that at 20 °C (Fig. 1b). Q₁₀ values of the peak current amplitude and the desensitization rate constant for the inward current remained below 2 across all temperature ranges, consistent with typical biological reactions and minimal temperature-dependence. Based on the outward current amplitude at 0 mV between 20 °C and 30 °C, the peak current Q₁₀ was 1.96 ± 0.18. In contrast, the sustained current showed a sharp increase in temperature sensitivity from 25 °C; based on the current amplitudes at 0 mV (20–30 °C), the sustained current Q₁₀ was 13.1 ± 1.1 (Fig. 4i), indicating strong thermosensitivity. To determine the thermal activation threshold for sustained current activation, we continuously recorded *Bma*-AVR-14B currents in the presence of 1 mM L-glutamate using a controlled temperature ramp (10 to 35 °C). The sustained current was activated above 24°C, with a mean threshold of 24.1 ± 0.6 °C (Figs. 1c, d). These results demonstrate that the *Bma*-AVR-14B possesses a unique thermosensitive property, characterized by the prominent activation of a large sustained current above 24 °C.

**Figure 1.**
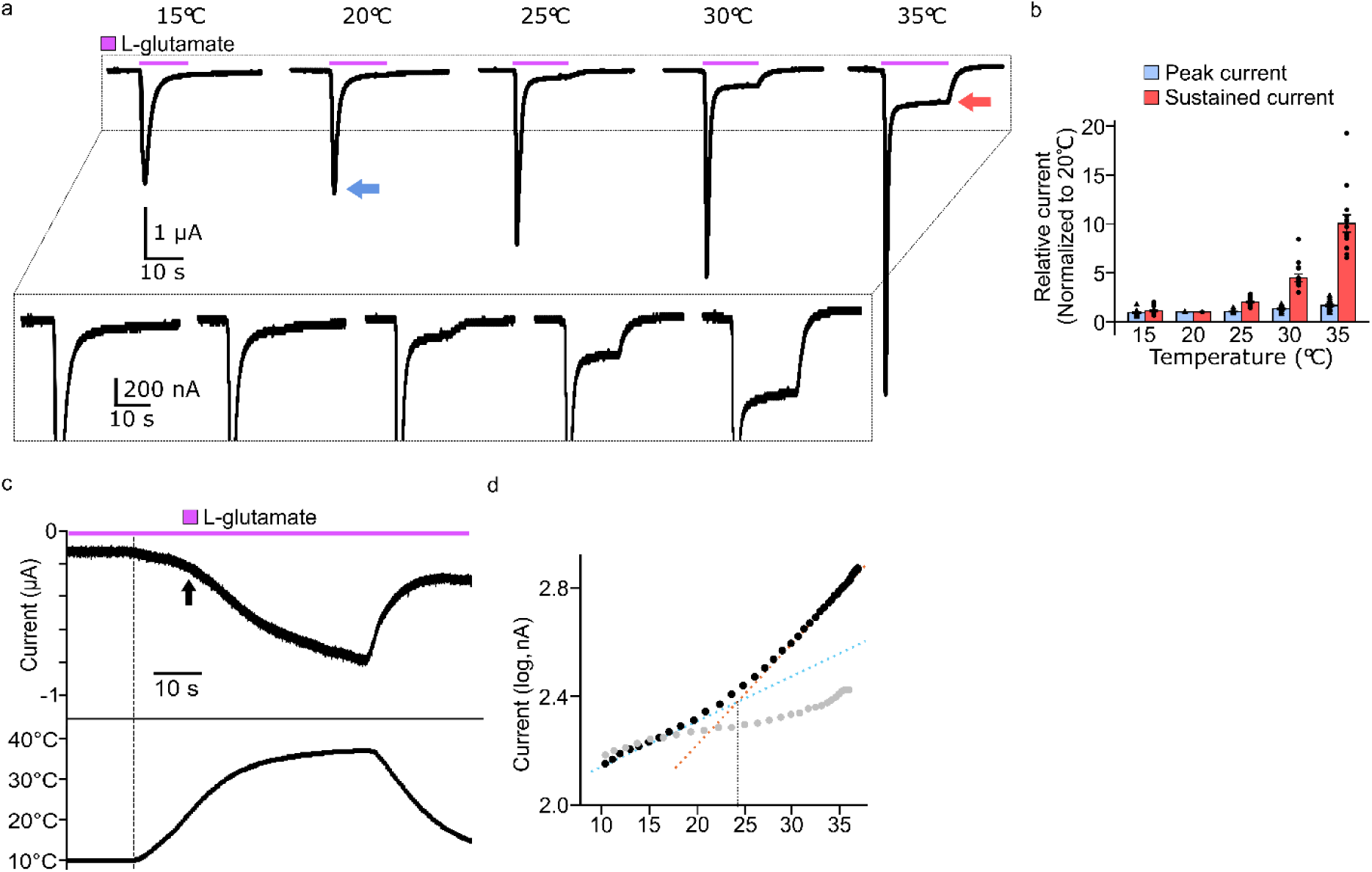
Biphasic current components and their modulation by temperature in *Bma*-AVR-14B. **a,** Representative L-glutamate–evoked currents recorded at –80 mV. Recording temperatures are indicated above each trace. Blue and red arrows mark the peak and sustained currents, respectively. The sustained current component is highlighted (inset, expanded view). **b**, Quantification of L-glutamate–evoked peak and sustained current amplitudes at each temperature, normalized to their respective values at 20 °C. Bars indicate means ± s.e.m., and dots denote individual biological replicates (n = 12). **c**, Representative current trace recorded during a continuous temperature ramp from approximately 10 °C to 35 °C (n = 9; upper). The actual recording temperature is shown in the lower trace. A vertical dotted line marks the onset of the temperature increase from the 10 °C baseline. **d**, Representative current–temperature relationships for *Bma*-AVR-14B (black) corresponding to (**c**) and for non-injected oocytes (gray). Linear fits to the baseline (blue) and activated (orange) regions were used to determine the activation temperature at their intersection.

### Characteristic of the temperature-dependent gating

Because the thermosensitive property of *Bma*-AVR-14B could arise from temperature-dependent ligand binding, we analyzed the concentration–response relationship for L-glutamate. The EC₅₀ values for L-glutamate did not differ significantly between 20 °C and 30 °C for either peak or sustained current components. At −80 mV, EC₅₀ values for the peak current were 0.51 ± 0.05 µM at 20 °C and 0.33 ± 0.05 µM at 30 °C, while those for the sustained current were 0.02 ± 0.01 µM at 20 °C and 0.05 ± 0.01 µM at 30 °C (Figs. 2a, b and Extended Data Fig. 1). A similar temperature-independence was observed at +20 mV under outward current conditions (Extended Data Fig. 1). These results indicate that the thermosensitivity of *Bma*-AVR-14B is not attributable to temperature-dependent changes in ligand binding. We also observed that the sustained current component was activated at lower concentrations of L-glutamate than the peak current, suggesting that distinct activation mechanisms underlie the two current components.

**Figure 2.**
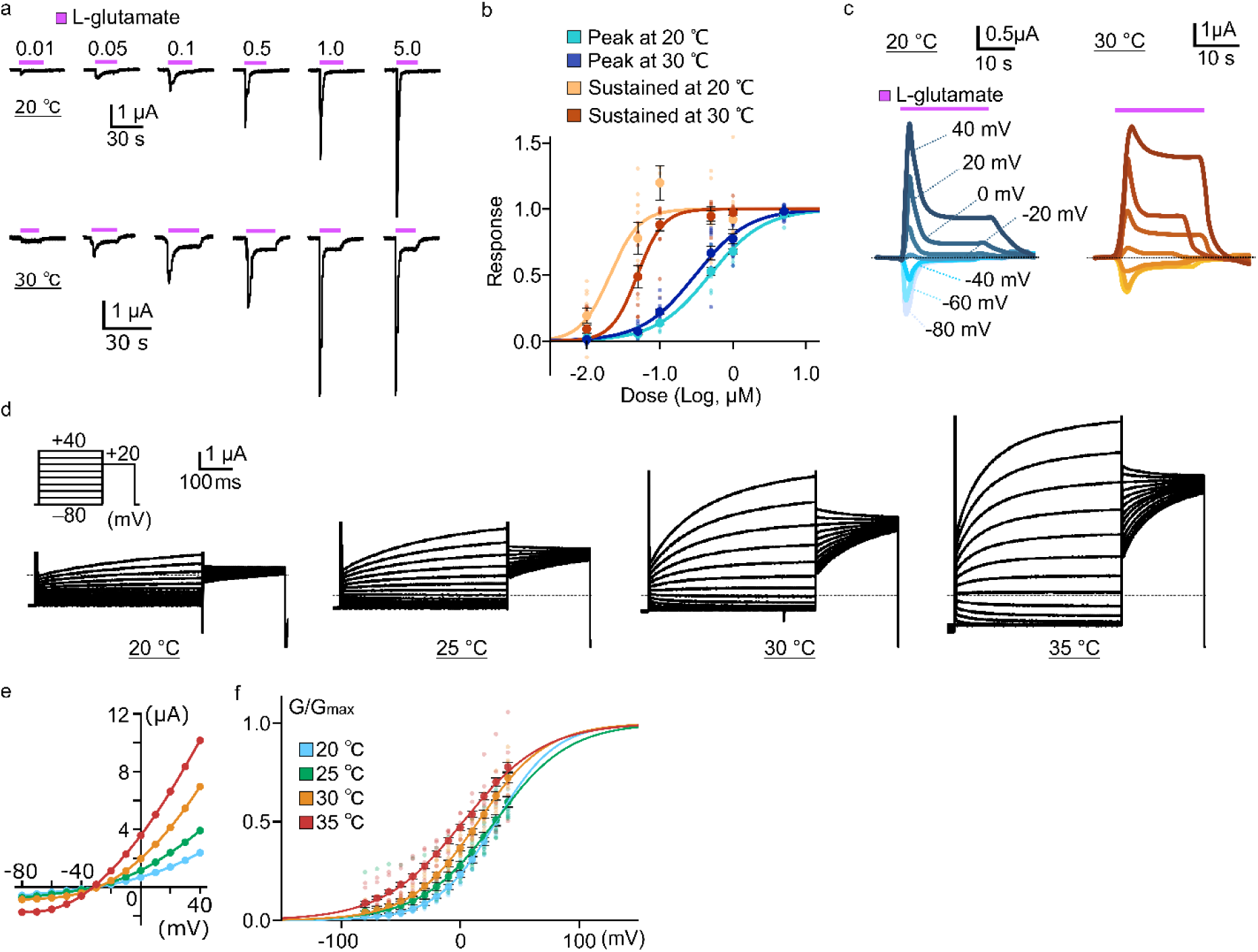
Temperature- and voltage-dependent gating of *Bma*-AVR-14B. **a**, Current traces evoked by the indicated concentrations of L-glutamate at −80 mV from the same oocyte. Recordings at 20 °C (top) and 30 °C (bottom) are shown. **b**, Normalized concentration–response relationships for peak and sustained currents at –80 mV (20 °C: n = 9; 30 °C: n = 11). **c**, Overlay of *Bma*-AVR-14B traces elicited by 1 mM L-glutamate, recorded from the same oocyte at holding potentials from −80 mV to +40 mV in 20 mV increments. Recordings at 20 °C (left) and 30 °C (right) are shown. **d**, Representative currents evoked by voltage step pulses applied after the sustained current reached a steady-state following the application of 1 mM L-glutamate. The pulse protocol, applied in 10 mV increments from −80 mV to +40 mV, is shown. Traces recorded from the same oocyte at 20 °C, 25 °C, 30 °C, and 35 °C are provided, with the dashed line indicating the baseline (0 µA). **e,** Representative current–voltage (I–V) relationships derived from the recordings shown in (**d**). **f,** Normalized conductance–voltage (G–V) relationships obtained from tail currents at +20 mV (20 °C: n = 10; 25 °C: n = 11; 30 °C: n = 11; 35 °C: n = 6). Recordings acquired from the same oocyte across the temperature set (20, 25, 30, and 35 °C) were treated as a single dataset. Bars indicate means ± s.e.m., and dots denote individual biological replicates.

We next analyzed L-glutamate–evoked currents of *Bma*-AVR-14B at various holding potentials. The sustained current exhibited a markedly larger amplitude at depolarized membrane potentials, and its amplitude was greater at 30 °C than at 20 °C (Fig. 2c), indicating that *Bma*-AVR-14B exhibits clear voltage-dependence and thermosensitivity. To further characterize the sustained current, voltage step protocols were applied after desensitization following L-glutamate application. From a holding potential of –80 mV, currents were recorded in response to the step-pulse protocol shown in Fig. 2d. The recordings revealed voltage-dependent changes during both the activation and tail phases, with current amplitudes markedly increased and kinetics accelerated at higher temperatures (Fig. 2d). The corresponding current–voltage (I–V) relationships showed a reversal potential near –30 mV, consistent with Cl⁻ selectivity under these conditions (Fig. 2e). Conductance–voltage (G–V) relationships obtained from tail currents exhibited voltage-dependent gating, showing a leftward shift of the activation midpoint (V₁/₂): 30.5 ± 1.9 mV at 20 °C; 32.8 ± 2.6 mV at 25 °C; 16.1 ± 1.8 mV at 30 °C; 4.4 ± 3.3 mV at 35 °C (Fig. 2f). This temperature-dependent shift in the G–V relationship has also been reported in TRP channels^23^. We concluded that the thermosensitivity of *Bma*-AVR-14B originates primarily from temperature-dependent gating modulated by voltage rather than changes in ligand binding.

### Pharmacological dissection of thermosensitivity

We next examined the pharmacological properties of *Bma*-AVR-14B using two representative modulators, ivermectin and thymol. Ivermectin is a widely used anthelmintic that targets pentameric ligand-gated ion channels in invertebrates and is reported to activate *Bma*-AVR-14B directly in the absence of L-glutamate, producing irreversible current^9,22^. We continuously recorded currents evoked by 10 nM ivermectin while gradually increasing the recording temperature from approximately 10 to 35 °C (Figs. 3a, b). Ivermectin-evoked currents were almost completely suppressed at low temperatures, and the currents exhibited pronounced thermosensitive activation, with thresholds around 23.0 ± 0.8 °C—closely matching those observed for L-glutamate–evoked sustained currents. These findings demonstrate that temperature is a critical determinant of ivermectin action on *Bma*-AVR-14B, revealing that pharmacological activation of GluCls is fundamentally temperature-modulated. Furthermore, the close match in thermal activation thresholds between the ivermectin-evoked and sustained currents suggest that both share a common gating mechanism.

**Figure 3.**
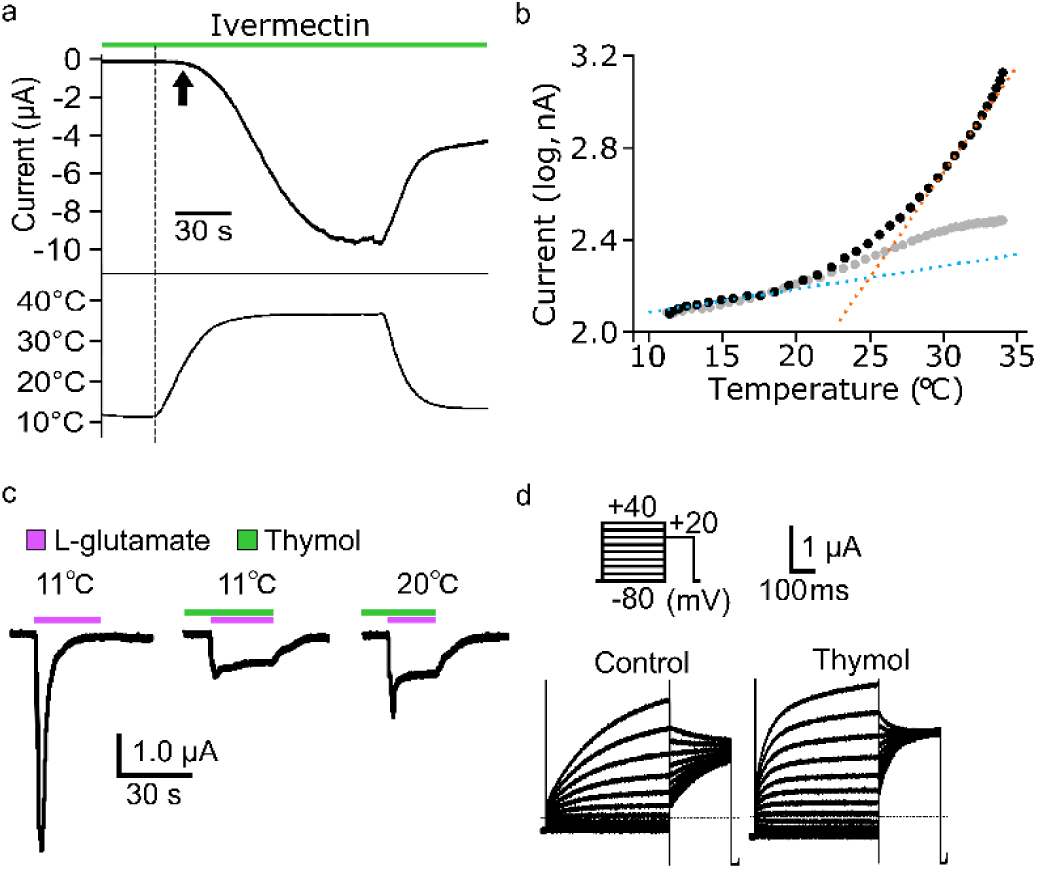
Pharmacological modulation of thermosensitivity in *Bma*-AVR-14B. **a**, Representative IVM–evoked current recorded at –80 mV during a continuous temperature ramp from approximately 10 °C to 35 °C (n = 12; upper). The actual recording temperature is shown in the lower trace. **b**, Representative current–temperature relationships obtained from (**a**). The intersections of the regression lines indicate the temperature thresholds. **c**, Representative L-glutamate–evoked currents recorded at 11 °C and 20 °C at a holding potential of –80 mV in the presence of thymol (11 °C: n = 11; 20 °C: n = 8). Current traces were obtained sequentially from the same oocyte to allow direct comparison across the indicated conditions. **d**, Representative current traces evoked by the voltage step protocol under control and thymol-treated conditions at 30 °C (n = 9), recorded from the same oocyte after application of 1 mM L-glutamate, as in Fig. 2d.

Thymol, a major constituent of thyme and oregano essential oils, is known for its insecticidal and antimicrobial properties and has been reported to block or modulate pentameric ligand-gated ion channel^24-26^. To examine the effect of thymol on the thermosensitivity of *Bma*-AVR-14B, we recorded L-glutamate–evoked currents at a holding potential of −80 mV in the presence or absence of thymol at 11, 20, and 30 °C. Thymol reduced the peak current amplitude (0.42 ± 0.18 at 11 °C, 0.53 ± 0.06 at 20 °C, and 0.68 ± 0.05 at 30 °C relative to control), while enhancing the sustained component (12.7 ± 1.6-fold at 11 °C, 5.7 ± 0.8-fold at 20 °C, and 2.5 ± 0.3-fold at 30 °C compared with control) (Fig. 3c and Extended Data Fig. 2). At temperatures below 20 °C, L-glutamate–evoked currents in the absence of thymol were negligible, indicating that thymol enables channel activation under those conditions where the wild-type channel is normally inactive. Although experimental limitations precluded precise determination of the thermal activation threshold below 11 °C, the presence of thymol enabled clear sustained currents to be evoked by L-glutamate even at 11 °C. These findings suggest that thymol lowers the thermal activation threshold of *Bma*-AVR-14B. To assess how thymol influences voltage-dependent gating, we applied voltage-step protocols to the same oocyte before and after thymol application at both 20 °C and 30 °C. The activation time constant (τ) decreased from 352 ± 41 ms to 94 ± 6 ms at 20 °C and from 246 ± 27 ms to 94 ± 10 ms at 30 °C, based on currents recorded at +40 mV (Fig. 3d). These results also indicate that thymol lowers the thermal activation threshold of the sustained component of *Bma*-AVR-14B.

### Structural basis of thermosensitivity

In this study, we identified two distinct current components in *Bma*-AVR-14B—a peak component and a sustained component—each exhibiting different EC₅₀ values for L-glutamate–evoked activation (Figs. 2a, b). Moreover, the two components responded differentially to pharmacological modulators (Fig. 3), suggesting that they arise from mechanistically distinct activation processes. Previous work based on the molecular dynamics simulation on the human glycine receptor α1 subunit (hGlyRα1), a pentameric ligand-gated ion channel with high homology to GluCl, has shown that, in addition to the canonical central axial pathway of the extracellular domain Cl⁻ can also enter through lateral fenestrations, forming a functionally distinct lateral pathways^16^. These computational insights raised the possibility that the two current components of *Bma*-AVR-14B in this study originate from physically and mechanistically separable permeation pathways. We therefore hypothesized that the thermosensitive sustained component arises from a permeation pathway distinct from that mediating the peak component. To test this hypothesis, we engineered *Bma*-AVR-14B mutants at conserved residues within the lateral pathway and characterized their current responses.

We generated an AlphaFold-predicted structural model of *Bma*-AVR-14B (Fig. 4a), which revealed two ion-access pathways: an axial pathway located in the center of extracellular domain (Fig. 4a, blue arrow) and five lateral pathways located at the subunit interfaces (Fig. 4a, red arrow). The residues lining these lateral pathways are conserved between *Bma*-AVR-14B and hGlyRα1 (Extended Data Fig. 3), suggesting a shared permeation mechanism. We focused on two highly conserved serine residues, S65 and S165, which form the entrance of the lateral pathway in the structural model (Figs. 4b–d) and correspond to hGlyRα1 S47 and A137, previously shown to contribute to ion permeation through the lateral pathways^16^ (Extended Data Fig. 3). Each residue was mutated to phenylalanine to constrict the permeation pathway or to glycine to enlarge it (Fig. 4d), and the mutant channels were electrophysiologically analyzed. Currents evoked by 1 mM L-glutamate were recorded from the same oocyte at both 20 °C and 30 °C (Figs. 4e–i and Extended Data Fig. 4). S165F nearly abolished the sustained current at both temperatures while preserving the peak current, indicating a selective loss of the temperature-dependent component (Figs. 4e, h). In contrast, single glycine substitutions produced currents comparable to wild-type, whereas the double S65G/S165G mutation selectively enhanced the sustained current at 0 mV at both temperatures (Figs. 4e, h). The double mutation also reduced the sustained current Q₁₀ value (Fig. 4i), consistent with increased permeation through an expanded lateral fenestration.

**Figure 4.**
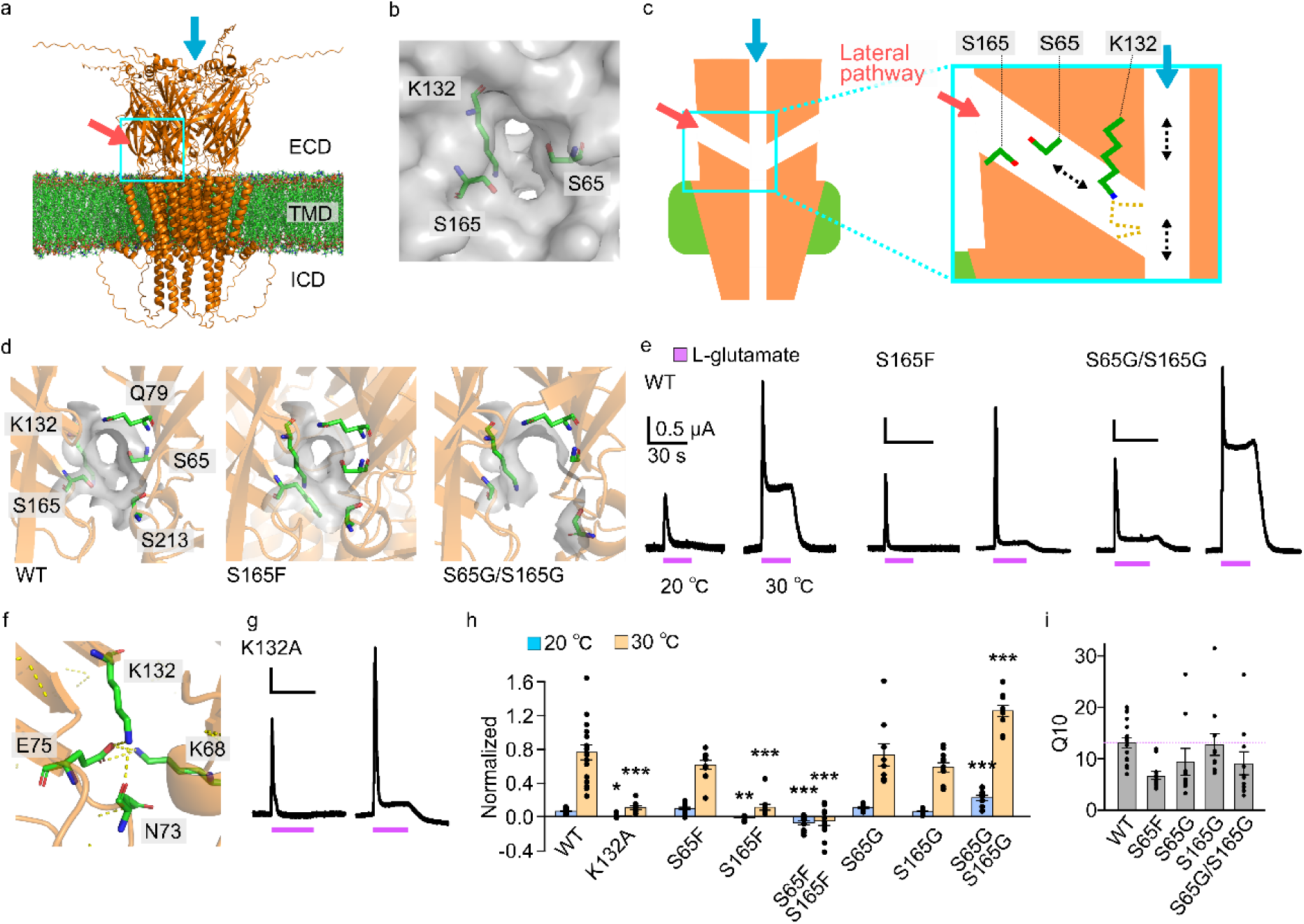
Structure–function coupling underlying temperature-dependent activation of *Bma*-AVR-14B. **a**, AlphaFold-predicted structural model of *Bma*-AVR-14B in a membrane. The central apical pore is highlighted by the red arrow, and the putative lateral ion-access route is indicated by the blue arrow. Protein backbone, orange; membrane, red headgroups/green acyl chains. **b**, An enlarged view of the lateral fenestration. Side chains forming the lateral pathway are shown as sticks. The molecular surface (gray) reveals a cavity consistent with the lateral opening. The view corresponds to the cyan arrow in (**a**). **c,** Schematic drawing of the lateral ion-entry route. **d**, Enlarged view of the lateral fenestration entrance, highlighting a minimal set of five residues that define the portal for the wild-type and the S165F and S65G/S165G mutants. Side chains are shown as sticks with per-residue molecular surfaces overlaid in gray. **e**, Representative 1 mM L-glutamate–evoked currents recorded at 0 mV at 20 °C and 30 °C. **f**, Deeper region of the fenestration showing the hydrogen-bond network (yellow dashed lines) among K132, E75, N73, and K68. **g**, Representative currents recorded from the K132A mutant at 20 °C and 30 °C. **h**, Normalized sustained currents for wild-type and mutants, expressed relative to the peak current recorded at 20 °C for each oocyte (17, 10, 11, 11, 12, 9, 11, 10; left to right). P values were analyzed using one-way ANOVA followed by Dunnett’s test versus wild-type (*P < 0.05; **P < 0.01; ***P < 0.001). **i**, The sustained current Q₁₀ values for wild-type and the mutants, with only mutants that exhibited detectable sustained currents included in the analysis (17, 11, 9, 11, 10; left to right). The magenta dashed line is provided for comparison with the WT. Bars indicate means ± s.e.m., dots denote individual biological replicates.

AlphaFold-guided modeling further identified a conserved hydrogen-bond network involving K132 and E75 deeper within the lateral pathway, in which K132 interacts with E75 and N73, and E75 contacts K68 from an adjacent subunit (Fig. 4f). To assess their functional contribution, we introduced charge-neutralizing mutations for conserved residues (K132A and E75S) to disrupt the network (Extended Data Fig. 3). K132A abolished thermosensitive sustained current without reducing the peak response (Figs. 4g, h), whereas E75S eliminated all detectable currents. Together, these results demonstrate that the lateral fenestration—spanning its entrance residues and the deeper K132-centered network—constitutes an integrated permeation pathway that is essential for generating the thermosensitive sustained current in *Bma*-AVR-14B. These mutations demonstrate that the lateral pathway forms a physically discrete and thermosensitive permeation route, distinct from the canonical axial pathway.

### Conserved thermal responsiveness of AVR-14B and its organism-level relevance

To assess whether thermosensitivity of anionic pentameric ligand-gated ion channel extends beyond parasitic nematodes, we first examined hGlyRα1. hGlyRα1 has been reported to lack voltage-dependence^27,28^, and application of 1 mM glycine at a holding potential of −80 mV evoked inward currents that exhibited desensitization followed by a sustained component. Recordings obtained at 12, 20, and 30 °C revealed an increase in the sustained component (Figs. 5a, b), and Q₁₀ analysis demonstrated that the dominant temperature-dependence occurs within the 12–20 °C range (Q₁₀ = 4.2 ± 0.4). Thus, these findings indicate that hGlyRα1 retains a detectable thermosensitivity in a sub-physiological range, raising the possibility of a conserved latent thermal responsiveness across species. We next evaluated the thermosensitivity of *C. elegans* AVR-14B using the temperature-ramp protocol as in Fig. 1c. *C. elegans* AVR-14B exhibited a thermosensitive sustained current comparable to that of *Bma*-AVR-14B (Figs. 5c, d). The temperature threshold was 22.3 ± 0.4 °C and the Q₁₀ of the inward sustained current amplitude analyzed at –80mV was 5.0 ± 0.6. These results suggest that thermosensitive sustained conduction through GluCls is evolutionarily conserved from nematodes to mammals.

**Figure 5.**
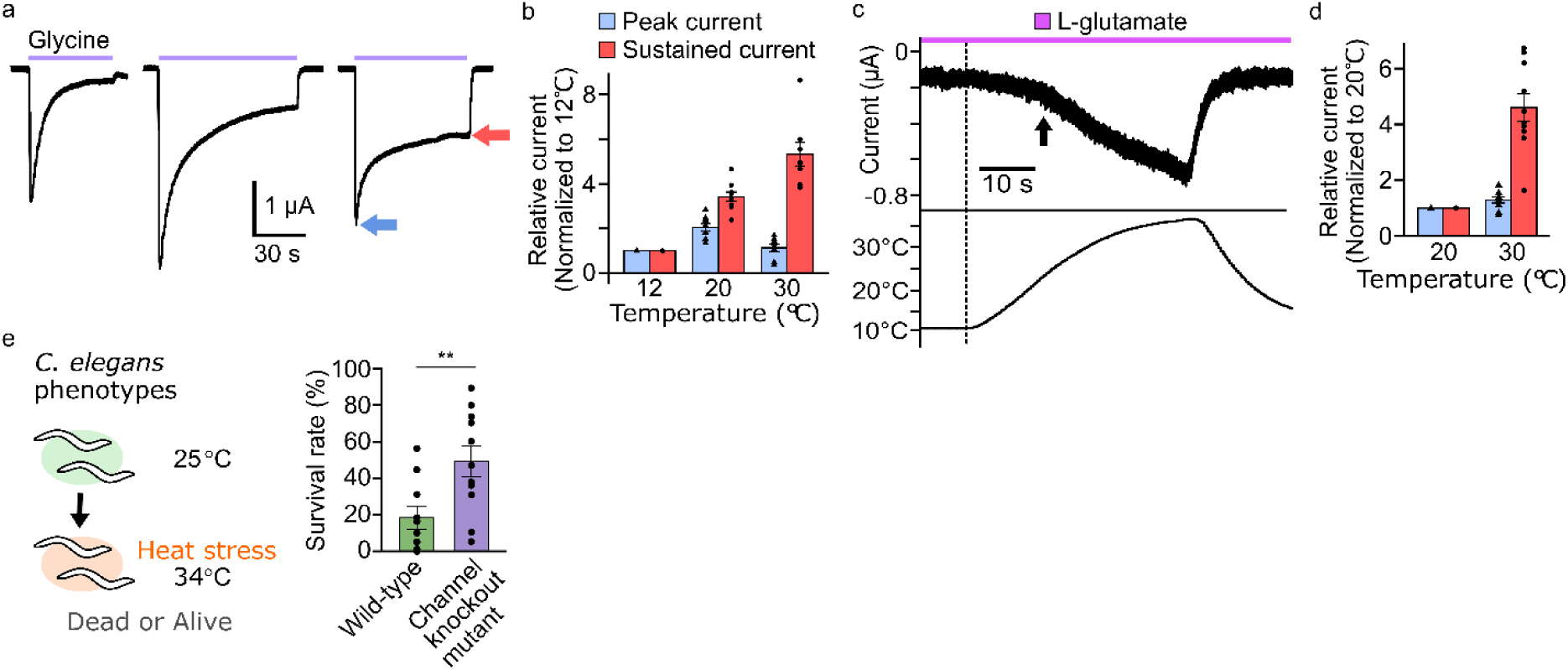
Comparative temperature dependence among species and potential physiological implications. **a,** Representative traces recorded at −80 mV for human glycine receptor α1 (hGlyRα1). Currents were evoked by 1 mM glycine and recorded at 12 °C, 22 °C, and 32 °C. Blue and red arrows indicate peak and sustained currents, respectively. **b**, Quantification of glycine–evoked peak and sustained current amplitudes from hGlyRα1 at each temperature, normalized to their respective values at 12 °C (20 °C: n = 10; 30 °C: n = 11). **c**, Representative current traces of *C. elegans* AVR-14B recorded at −80 mV during a continuous temperature ramp from approximately 10 °C to 35 °C in the presence of 1 mM L-glutamate (n = 8). **d**, Quantification of glycine–evoked peak and sustained current amplitudes from *C. elegans* AVR-14B, normalized to their respective values at 20 °C (n = 10). **e**, Heat tolerance assay in *C. elegans*, comparing wild-type animals and *C. elegans* AVR-14 channel knockout mutants (wild-type: n = 10; mutant: n = 11). Animals were raised at 25 °C until adulthood and then exposed to 34 °C for 24 h. P values were analyzed using *t*-test (**P < 0.01). Bars indicate means ± s.e.m., and dots denote individual biological replicates.

The activation threshold of the thermosensitive sustained current in AVR-14B (approximately 24 °C) closely matches the upper limit of the optimal temperature range for *C. elegans* growth (∼25 °C)^29^. Moreover, since ivermectin exerts its lethal effect by constitutively activating GluCls^30^, including AVR-14B, we reasoned that temperature-dependent sustained activation of AVR-14B might similarly compromise nematode survival under heat stress. To test whether this excessive channel activity is detrimental and whether the temperature-dependence of AVR-14B has organism-level physiological significance, we assessed the heat tolerance of an *avr-14*–null strain. The knockout *C. elegans* displayed significantly improved survival relative to wild-type controls (wild-type: 18.3 ± 7.0%, *avr-14*–null: 49.2 ± 8.4%; Fig. 5e), indicating that excessive chloride influx through AVR-14B’s thermosensitive sustained currents contributes to the detrimental consequences of prolonged heat exposure.

## Discussion

Here, we demonstrate intrinsic thermosensitive gating in GluCl inhibitory channels. This property suggests that temperature can modulate inhibitory ligand-gated ion channels across multiple levels, from channel architecture to physiological signaling and pharmacological responsiveness.

In this study, we found that glutamate-evoked *Bma*-AVR-14B currents exhibited a biphasic profile, comprising a rapidly desensitizing peak component and a non-desensitizing sustained component, and that the sustained component was temperature-sensitive. Mechanistically, the ligand-evoked responses did not show temperature-dependent changes, which is distinct from AMPA-type receptors where temperature-dependent ligand recognition contributes to the emergence of thermosensitivity^31,32^. By contrast, the *Bma*-AVR-14B sustained current showed weak voltage-dependence, and its G–V curve shifted leftward with increasing temperature. This temperature-driven facilitation of activation resembles the voltage and temperature sensitivity observed in thermo-TRP channels, which integrate both temperature and voltage gating into their activation mechanisms^23^.

Thermosensitivity was shown here to be closely associated with ion currents permeating through the lateral pore and its gating. Although lateral pores, including vestigial traces, are structurally conserved among Cys-loop receptors, their functional contribution to permeation differs fundamentally across receptor classes. Structural comparisons reveal a clear dichotomy: in cation-selective nicotinic acetylcholine receptors (Extended Data Fig. 5a), the negatively charged axial pore provides the dominant conductive pathway, whereas lateral pores are not sufficiently wide to accommodate ion permeation^15^. In anion-selective GlyRα1 (Extended Data Fig. 5b), the axial pore is unfavorable for anions, but the lateral pore is wide and positively charged, suggesting that lateral conduction is possible, as indicated by simulation studies^16^. Consistent with this anion-selective architecture, the AlphaFold model of *Bma*-AVR-14B exhibits a negatively biased axial pore alongside an open, positively charged lateral pore (Extended Data Fig. 5c). Under such conditions, sustained ion permeation through the axial pore may be disfavored. Together, these examples reveal that Cys-loop receptors employ diverse strategies to utilize lateral pathways, and that the structural diversity of lateral pores may underlie conformational flexibility that enables temperature-dependent changes in ion conduction.

The lateral pore–mediated sustained current may also contribute to the temperature-dependence of ivermectin action. Ivermectin-evoked GluCl currents showed strong temperature-dependence, with a threshold near 24 °C and little activity at about 10 °C, closely matching the thermal profile of the L-glutamate-evoked sustained current. These shared properties suggest that ivermectin may engage the same temperature-dependent gating mechanism that underlies the sustained current. Structural studies place the ivermectin-binding pocket at the extracellular membrane interface of the transmembrane domain, in close proximity to the lateral pore, providing a functional framework for probing channel activation mechanisms. This membrane-adjacent region also serves as a hotspot for lipophilic modulators of Cys-loop receptors^33-35^, highlighting its importance in shaping receptor channel behavior. These mechanistic features have direct implications for how ivermectin acts *in vivo* under changing thermal conditions. Avermectins, including ivermectin, are widely used antiparasitic agents and agricultural pesticides that act on GluCl channels in invertebrates. Previous studies reported that ivermectin efficacy is markedly reduced at low temperature, such as 5 °C in *C. elegans*, and this reduction was primarily attributed to impaired uptake or slowed biochemical processes^36^. Our findings instead demonstrate that the primary molecular target itself is temperature-sensitive, identifying temperature as an overlooked pharmacological parameter in avermectin action. This insight calls for a reassessment of ivermectin’s physiological effects across thermal conditions, beyond explanations based solely on uptake, metabolism, or organismal activity. In addition, target thermosensitivity may alter the strength of toxic effects on other organisms in the environment under different thermal conditions^37-39^.

The activation threshold of AVR-14 (∼24 °C) matches the upper limit of the growth temperature range of nematodes. Intrinsic thermosensitivity of parasitic GluCl implies that inhibitory chloride channels contribute to thermal adaptation during vector-to-host transmission, when parasites experience steep thermal gradients upon entering vertebrate hosts (> 25 °C). In *C. elegans*, temperatures above 25 °C are known to be detrimental, inducing marked changes in metabolism and behavior^29^. *C. elegans* AVR-14 functions as an inhibitory component in neural circuits, stabilizing behavioral output such as reversal-based avoidance^40-42^. Its thermosensitivity indicates that temperature responsiveness can arise within neural circuits not classically considered temperature-dependent, providing a plausible neural pathway through which temperature can influence behavior. Consistent with this idea, *avr-14* is broadly expressed across diverse neuron types^43^, including thermosensory neurons, suggesting that GluCl-mediated temperature-dependent modulation extends beyond interneurons. Several sensory neurons co-express AVR-14 together with established various thermosensors^4,44,45^, raising the possibility that GluCls act alongside other thermosensors to shape thermal responses within neural circuits. Moreover, our finding that AVR-14 contributes to heat tolerance reveals a neural component to this phenotype, which has traditionally been attributed to metabolic and transcriptional stress pathways^46^.

In addition to the phasic (peak) current, we identified a sustained current in GluCl that is thermosensitive, has been largely overlooked, and whose physiological significance remains unclear. The two-component property in GluCl is reminiscent of the two canonical modes of Cys-loop–mediated inhibition—phasic inhibition mediated by desensitizing peak currents and tonic inhibition mediated by non-desensitizing sustained conductance. In this framework, the lower EC50 of the sustained component in *Bma*-AVR-14B is consistent with the emergence of tonic inhibition at lower agonist concentrations (Figs. 2a, b). In general, phasic inhibition refines fast synaptic signaling on short timescales, whereas tonic inhibition provides a continuous inhibitory conductance that sets the baseline level of excitability and is implicated in neuronal development, sensory processing and state regulation^47,48^. These two modes are typically shaped by differences in subunit assembly, paralog composition, subcellular localization, and neurotransmitter time course. Our findings demonstrate that these two functionally distinct modes can coexist within a single GluCl as a separable current component. Conservation of this property suggests that temperature can bias the balance between phasic and tonic inhibition, revealing an unrecognized dimension of inhibitory control. Because glycinergic inhibition controls sensory gain^8^, temperature-dependent modulation of GlyRs may influence inhibitory tone in inflammatory pain and during hypothermia or therapeutic cooling. Future work combining in vivo electrophysiology, behavioral analyses, and mammalian models will be required to determine how inhibitory thermosensation is integrated into thermal processing in intact nervous systems.

## Methods

### Molecular biology

*B. malayi avr-14B*, *C. elegans avr-14B*, and human GLRA1 gene fragments were obtained from eBlocks Gene Fragments and subsequently cloned via ligation into the pGEMHE vector. Point mutations were generated using the PrimeSTAR Mutagenesis Basal Kit (TAKARA BIO) and validated through DNA sequencing. Complementary RNAs (cRNAs) were synthesized by transcription of linearized plasmids using the mMACHINE T7 Transcription Kit (Thermo Fisher Scientific Inc).

### Xenopus oocytes preparation

All studies were conducted following the ethical standards set by the Animal Experimentation Ethics Committee of Hiroshima University and conformed with generally agreed international regulations. Oocytes were surgically harvested from *Xenopus laevis* anesthetized by immersion in ice-cold water containing tricaine. The isolated oocytes were treated with type I collagenase (1.0 mg/ml, Sigma-Aldrich) for 40 minutes to 2 hours, injected with 50 nl of cRNA, and incubated at 18 °C in modified Barth’s Solution (MBSH: 88 mM NaCl, 1 mM KCl, 2.4 mM NaHCO_3_, 0.3 mM Ca (NO_3_)_2_, 0.41 mM CaCl_2_, 0.82 mM MgSO_4_, and 15 mM Hepes (pH 7.6)).

Incubation time was 2 to 3 days for AVR-14B, whereas for human GlyRα1, it was 6 to 16 h.

### Two-electro voltage-clamp

Recordings were performed from oocytes using a bath-clamp amplifier (OC-725D, Warner Co), an AD/DA converter, and pClamp 10 software (Molecular Devices)^49^. Glass microelectrodes filled with 2 M CH₃COOK and 1 M KCl (pH 7.2) were used as intracellular electrodes, with resistances ranging from 0.2 to 1.0 MΩ. The bath solution contained 98 mM NaCl, 2 mM KCl, 2 mM MgCl₂, and 5 mM Hepes (pH 7.4, adjusted with NaOH). Cl^−^ currents induced by L-glutamic acid and glycine administration were recorded at a holding potential of −80, 0 and +20 mV with the DC gain booster of the amplifier activated (Figs. 1, 2a−c, 3a−c, 4e, 4g and 5a, 5c and Extended Data Figs. 1a, 2, 3a, 3b). The voltage-step protocol was performed with the membrane potential held at −80 mV. Step pulses (500 ms) were applied from −80 to +40 mV in 10 mV increments, with a deactivating tail potential of +20 mV (Figs. 2d and 3c). Recordings were carried out under continuous perfusion using a VC3-8PG perfusion system (ALA Scientific Instruments, Inc). The temperature was controlled during perfusion using either warmed or ice-cold buffer^50^. After equilibration to the desired temperature, L-glutamic acid (FUJIFILM Wako Chemical; adjusted to pH 7.4) or glycine (Nacalai Tesque Inc), dissolved in temperature-equilibrated buffer, was applied and immediately mixed. For repetitive stimulation protocols, an inter-stimulus interval of ≥ 4 min was employed for glycine-induced activation of hGlyRα1 (Fig. 5a), whereas an interval of ≥ 1 min was used for glutamic acid-induced activation of worm AVR-14B channels (Figs. 1a, 2a, 3c, 4e and 4g). For temperature-threshold measurements, a continuous temperature ramp was generated by perfusing a ligand-containing warm solution (Figs. 1c, 3a and 5c). Thymol (Tokyo Chemical Industry, TCI) and ivermectin (Tokyo Chemical Industry, TCI) were dissolved in DMSO and diluted into the buffer to final concentrations of 300 μM and 10 nM, respectively, with a final DMSO concentration of 0.1% in buffer (Fig. 3). For control experiments without drugs, 0.1% DMSO was added to the buffer.

### Structural modeling / Visualization

The three-dimensional structural models are constructed by AlphaFold using the amino acid sequence^51^. Amino acid sequences of the channel proteins are obtained from NCBI (*Bma*-AVR-14B: GenBank: QRW38687.1; hGlyRα1: NCBI Reference Sequence: NP_001139512.1) and corresponding mutants. The *Bma*-AVR-14B protein was positioned within the membrane using the PPM server, and subsequently embedded in a lipid bilayer using CHARMM-GUI Membrane Builder. Molecular graphics and analyses were performed using the open-source version of PyMOL (The PyMOL Molecular Graphics System, Version 3.1.0, Schrödinger, LLC). Residues are shown as sticks with standard PyMOL elemental coloring. Van der Waals surfaces of the wild-type channel and mutants were generated without including water molecules and other solvent. Electrostatic surface potentials were calculated using the Adaptive Poisson–Boltzmann Solver (APBS). Structural models for *Bma*-AVR-14B were obtained from AlphaFold predictions, whereas experimentally determined atomic coordinates for other Cys-loop receptors were derived from published Protein Data Bank (PDB) structures, including the open-state nicotinic acetylcholine receptor (human nAChR: 7KOX) and the open-state glycine receptor (zebrafish GlyRα1: 6PM6)^52,53^. All structures were prepared via the PDB2PQR pipeline on the Poisson–Boltzmann web server (https://server.poissonboltzmann.org/).

Protonation states were assigned using PROPKA at pH 7.0, and partial charges and atomic radii were applied according to the AMBER force field. Electrostatic potential maps were computed by solving the linearized Poisson–Boltzmann equation with APBS under default ionic strength conditions. The resulting electrostatic potential maps were visualized on the molecular surface using the APBS plugin in PyMOL with a consistent color scale across all structures to facilitate direct comparison of charge distributions.

### Heat tolerance assay

The N2 (Bristol) wild-type strain, CX12709 *avr-14B (ad1302)* strain, and *E. coli* OP50 were provided by the CGC, which is funded by NIH Office of Research Infrastructure Programs (P40 OD010440). The strains were maintained following standard procedures^54^. *C. elegans* were cultured on nematode growth medium (NGM) containing 2% (w/v) agar with a lawn of *E. coli* OP50 in 60-mm dishes at 18 °C and transferred to fresh NGM every 3–4 days. *C*.

*elegans* has been widely used to study conserved responses to heat stress, with reported heat stress conditions ranging from 28 to 40°C depending on the laboratory, for worms that normally survive and reproduce between 12 and 26 °C^29,46,55^. Based on these previous studies, we assessed survival under conditions in which approximately 30–50% of wild-type worms are expected to alive: five to ten well-fed adult worms were placed on a 35-mm dish containing NGM for egg-laying for 1.5–2 h. The progeny of about 70 worms were cultured at 25°C for 62–75 h and incubated at 34 °C for 24 h.

### Sequence alignment and homology analysis

Pairwise sequence identity, similarity, and gap fraction between *Bma*-AVR-14B and hGlyRα1 were calculated using EMBOSS Needle (v6.6.0) with the BLOSUM62 substitution matrix. Multiple sequence alignment of *Bma*-AVR-14B, *Cel*-AVR-14B, and hGlyRα1 was performed using Clustal W (BLOSUM62).

### Electrophysiological data analysis

Response amplitudes were calculated by subtracting the baseline current, defined as the mean current over 0.5 seconds immediately before ligand application, from both the peak current (mean over 0.01 seconds at the maximum) and the sustained current (mean over 0.5 seconds during the sustained phase). The activation time constant was obtained by fitting the time course of current decay from the peak to the sustained level with a single-exponential function^49^. I(t) = I_sustained_ + A*exp (– (t–t_peak_)/τ). Fittings were performed on the peak-to-sustained segment by least-squares; the resulting τ value is reported for the dominant exponential component. For mutant analysis in Fig. 4, to account for differences in expression among mutants, sustained currents were normalized to the corresponding peak amplitude from the same trace and expressed as the sustained/peak ratio. Voltage step protocols were applied after the residual current had stabilized following the initial peak response to measure the L-glutamate–evoked sustained currents remaining after desensitization. Macroscopic and tail currents were quantified by averaging 0.5-ms windows at the end of the voltage step and immediately after repolarization, respectively. Conductance–voltage (G–V) relationships were obtained from tail currents, normalized within each cell, and fitted with a Boltzmann Z-delta model in Clampfit to estimate the V_1/2_^49^. The function is defined as f (V) = V_min_ + (V_max_ − V_min_) / (1 + exp ((zd * F / (R * T)) * (V − V_1/2_))), where V_min_ and V_max_ are the minimum and maximum potentials, respectively; zd is the effective valence, F is the Faraday constant, R is the gas constant, T is the temperature, and V_1/2_ is the midpoint potential. Figures of the fitted curves were generated in Python using the best-fit parameters obtained in Clampfit. The mean response amplitudes for each L-glutamate concentration were plotted and fitted with the Hill equation to determine the EC_50_ values using Clampfit 11 (Molecular Devices). Because the current induced by 5 mM L-glutamate was reduced compared to that at 1 mM, concentration–response curve fitting for sustained currents was limited to 1 mM to avoid these confounding effects. Data visualization and additional curve fitting were performed in Python (NumPy, SciPy, Matplotlib).

### Thermosensitivity analysis

The temperature coefficient Q₁₀ was calculated from response amplitude measured at two temperatures, T1 and T2 (°C), using the standard definition Q₁₀ = (I_2_/I_1_)^10/(T2−T1)^. Q₁₀ denotes the fold-change per 10 °C; values of ≈ 1 indicate negligible temperature dependence, ≈ 2 moderate dependence, and > 3 indicates strong temperature-dependence^56-58^. For desensitization, Q₁₀ was computed on the rate k = 1/τ, such that Q₁₀ > 1 indicates faster desensitization with increasing temperature (shorter τ), and Q₁₀ < 1 indicates slower desensitization (longer τ). Thermal thresholds for channel activation were determined as follows. Current data (nA) in the presence of glutamate or drug treatment were downsampled at 1-s intervals and log-transformed, then plotted against temperature (°C). Log transformation was used to linearize the exponential increase in current. The thermal threshold was defined as the intersection of two linear fits on either side of the inflection point in the log-current versus temperature plot.

### Statistical analysis

All data, except for Q_10_ values, are presented as means ± s.e.m, with individual recordings shown as overlaid dots in the graphs. Dots, individual oocytes. The number of samples (n) and error bars are indicated in the figure legends. Data were collected at least three independent batches of oocytes for electrophysiological recording or four separate days for *C. elegans* behavior assay. The sample sizes were determined established practices and previous studies^4,44^, and parametric statistical analyses were applied in line with prior work using comparable experimental designs. Single comparison in Fig. 5e was performed using unpaired Student’s *t*-test. Multiple comparisons were performed using one-way ANOVA followed by Tukey–Kramer’s post hoc test and Dunnett’s test. All statistical tests were two-sided. Analyses were conducted using EZR software^59^. Significance was defined as p < 0.05 (*p < 0.05, **p < 0.01, ***p < 0.001). No outlier removal was performed. Statistical analyses of EC₅₀ values were performed using log-transformed data to satisfy assumptions of normality.

## Data Availability

All data generated during this study are included in the manuscript. Source data are provided with this paper and includes raw data and statistical analysis data. The AlphaFold-predicted structural models generated and/or analyzed during this study are available from the corresponding author upon reasonable request.

## Acknowledgments

We thank the members of Physiology and Biophysics laboratory (Hiroshima University) for supports and discussions. DNA sequencing was performed using the facilities of the Natural Science Center for Basic Research and Development (N-BARD) at Hiroshima University [NBARD-TU4Z21VP], which are supported by the MEXT Program for supporting construction of core facilities (Grant Number JPMXS0441300026). K.O. is supported by a Grant-in-Aid for Young Scientists (KAKENHI 23K14235) and Grant-in-Aid for JSPS Fellows (KAKENHI 25KJ0244). Y.F. is supported by a research grant from the Nakatani Foundation.

## Author information

### Authors and Affiliations

Physiology and Biophysics, Graduate School of Biomedical and Health Sciences (Medical), Hiroshima university, Hiroshima, Japan.

Kohei Ohnishi & Yuichiro Fujiwara

## Contributions

K.O. performed experiments, data analysis, and contributed to study design and manuscript preparation. Y.F. supervised the project, contributed to study design, data interpretation, and manuscript writing.

---

**Extended Data Figure 1.**
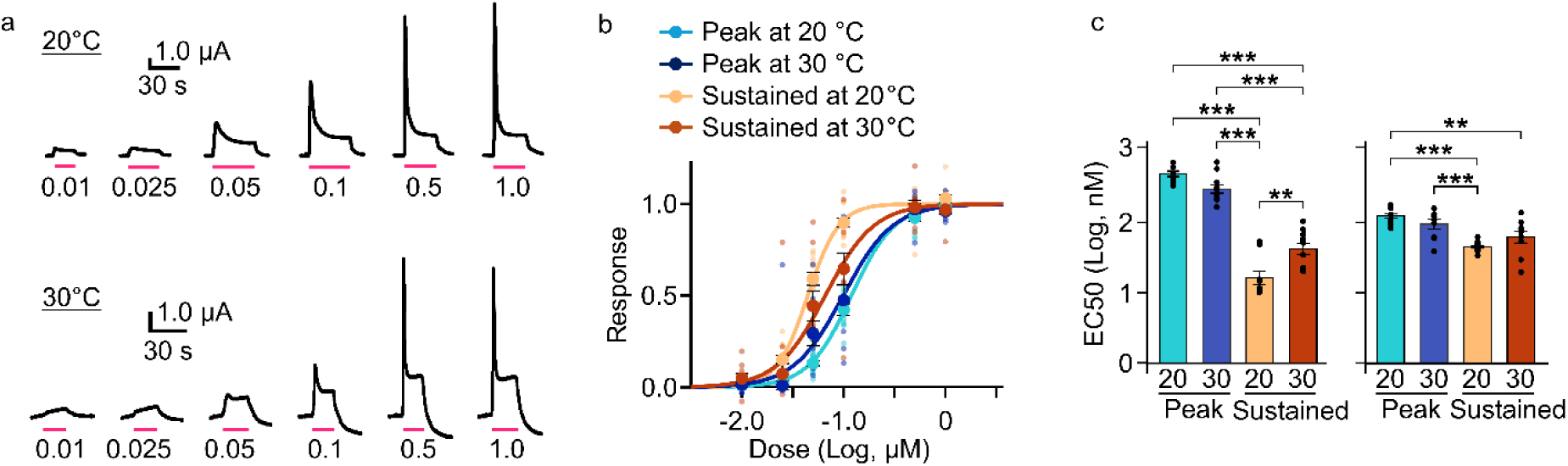
Dose–response relationship of L-glutamate. **a**, Representative current traces evoked by increasing concentrations of L-glutamate recorded at +20 mV at 20 °C and 30 °C. **b** Concentration–response relationships obtained by hill fittings for peak and sustained currents at +20 mV. **c**, Summary of EC_50_ from Hill fittings for peak and sustained currents at –80 mV and +20 mV at 20 °C and 30 °C (–80 mV, 20 °C: n = 9; –80 mV, 30 °C: n = 11; +20 mV, 20 °C: n = 12; +20 mV, 30 °C: n = 9). Bars indicate means ± s.e.m., and dots denote individual biological replicates. P values were analyzed using one-way ANOVA followed by Tukey-Kramer’s test (*P < 0.05; **P < 0.01; ***P < 0.001).

**Extended Data Figure 2.**
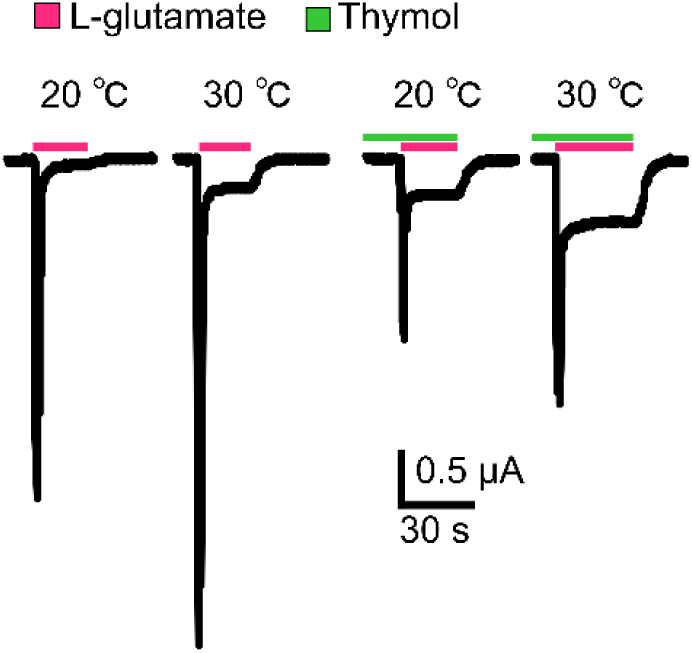
Representative L-glutamate–evoked currents recorded at 20 °C and 30 °C at a holding potential of –80 mV in the presence of thymol (20 °C: n = 14; 30 °C: n = 10). Current traces were obtained sequentially from the same oocyte to allow direct comparison across the indicated conditions.

**Extended Data Figure 3.**
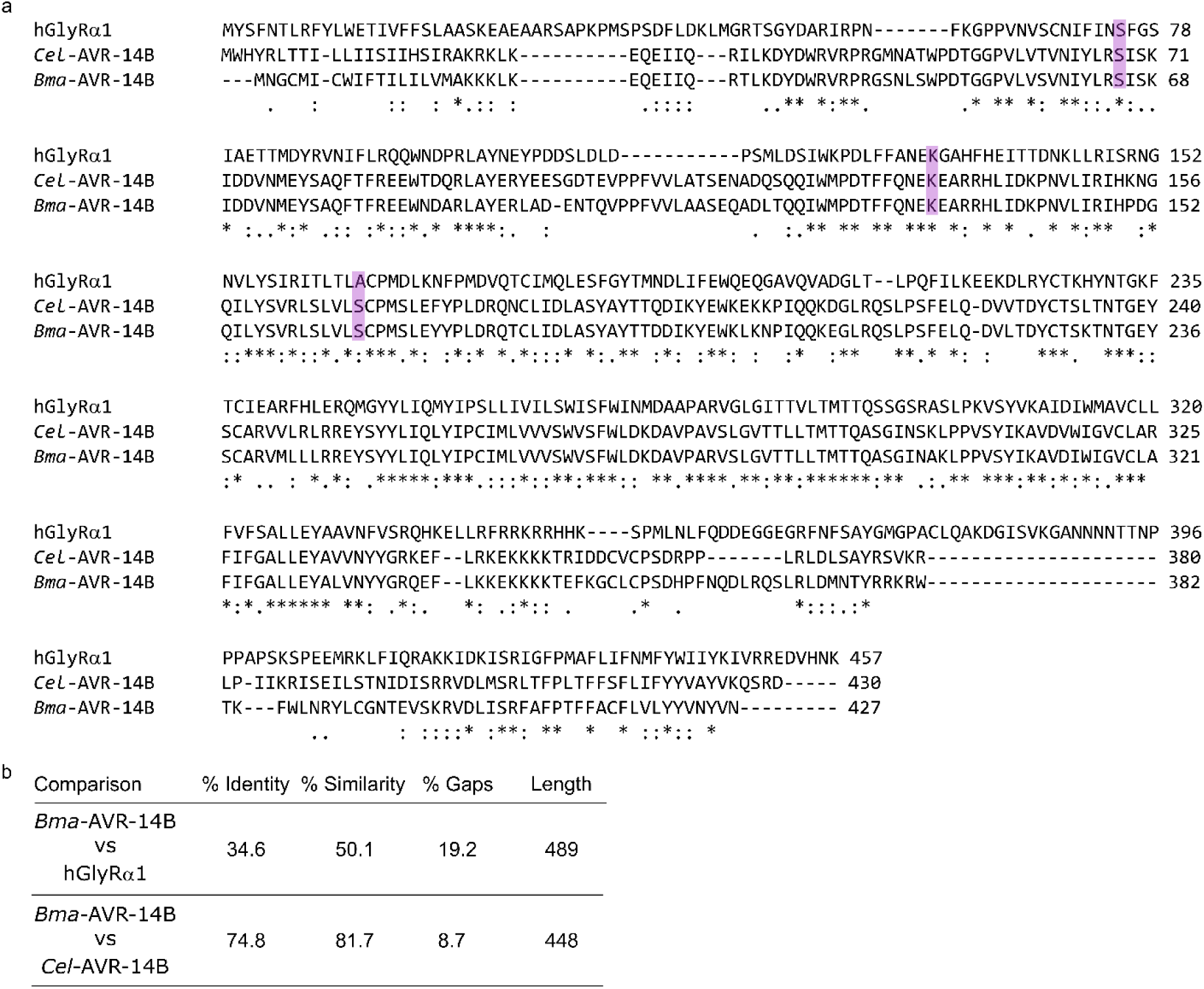
Sequence conservation of nematode AVR-14B and human GlyRα1. **a**, Multiple sequence alignment of *Bma*-AVR-14B, *Cel*-AVR-14B, and human GlyRα1 performed using Clustal W (BLOSUM62). Conserved positions are indicated by “\**”, “:”, and “.” according to Clustal notation (“*” = identity; “:” = strong conservation; “.” = weak conservation), and* residues targeted for mutagenesis are highlighted by magenta boxes. **b**, Pairwise sequence identity, similarity, and gap fraction between *Bma*-AVR-14B and human GlyRα1 calculated using EMBOSS Needle (BLOSUM62).

**Extended Data Figure 4.**
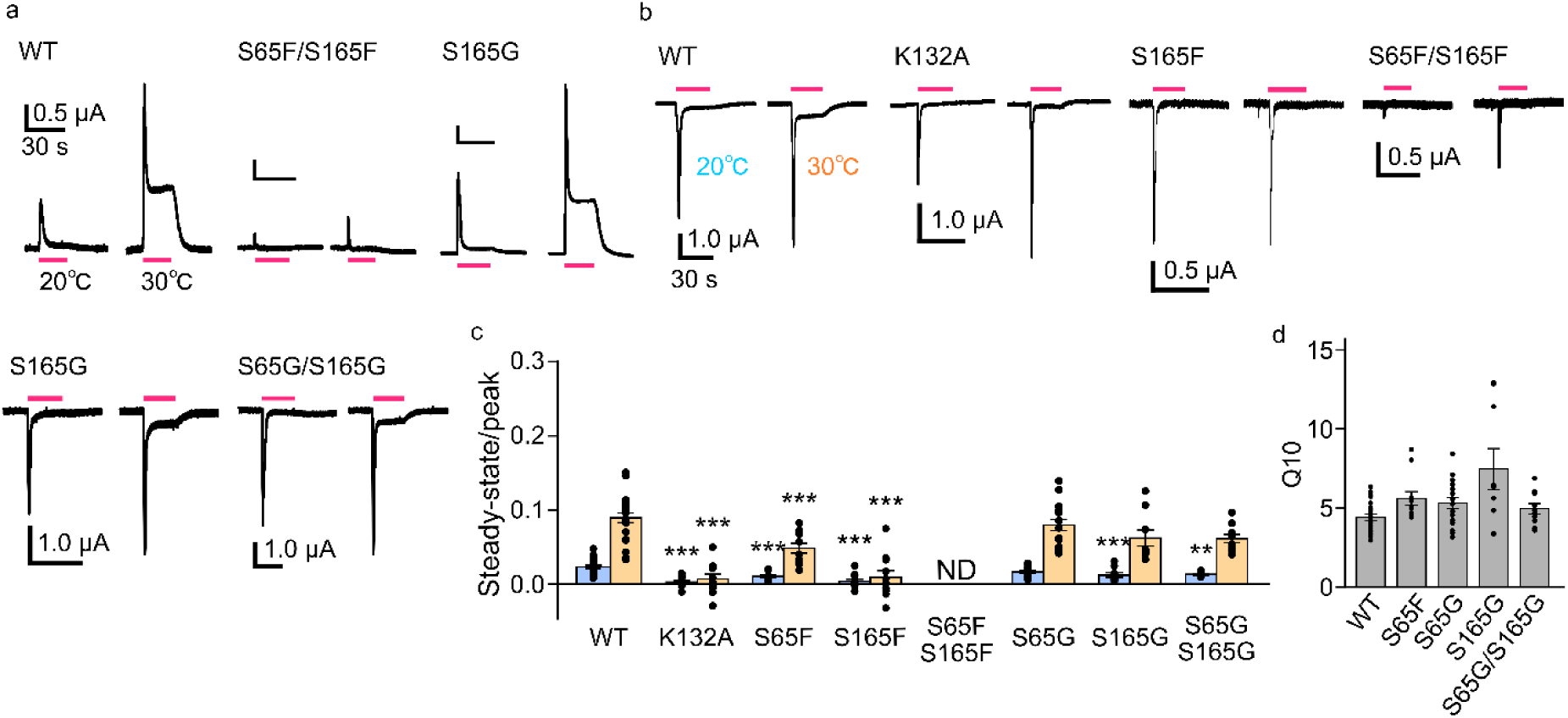
Electrophysiological characterization of wild-type and mutant channels. **a**, Representative L-glutamate–evoked currents recorded at 0 mV at 20 °C and 30 °C. The wild-type trace is identical to that shown in Figs. 4**e**. **b**, Representative L-glutamate–evoked currents recorded at −80 mV at 20 °C and 30 °C. **c**, Normalized sustained currents at 20 °C and 30 °C, expressed relative to the peak current measured at 20 °C for each recording (n = 23, 11, 11, 11, 9, 16, 9, and 10 from left to right). S65F showed wild-type–like currents at 0 mV but reduced inward currents at −80 mV. The S65F/S165F double mutant reduced both peak and sustained currents; because peak current amplitudes were substantially decreased, sustained current values were not determined (ND). Because the peak reduction is likely secondary to severe constriction of both lateral entrances rather than altered gating, this mutant was not used to draw mechanistic conclusions in the main text. P values were analyzed using one-way ANOVA followed by Dunnett’s test versus wild-type (***P < 0.001). **d**, The sustained current Q₁₀ values for wild-type and the mutants, with only mutants that exhibited detectable sustained currents included in the analysis (n = 23, 11, 16, 9, and 10 from left to right). Bars indicate means ± s.e.m., and dots denote individual biological replicates.

**Extended Data Figure 5.**
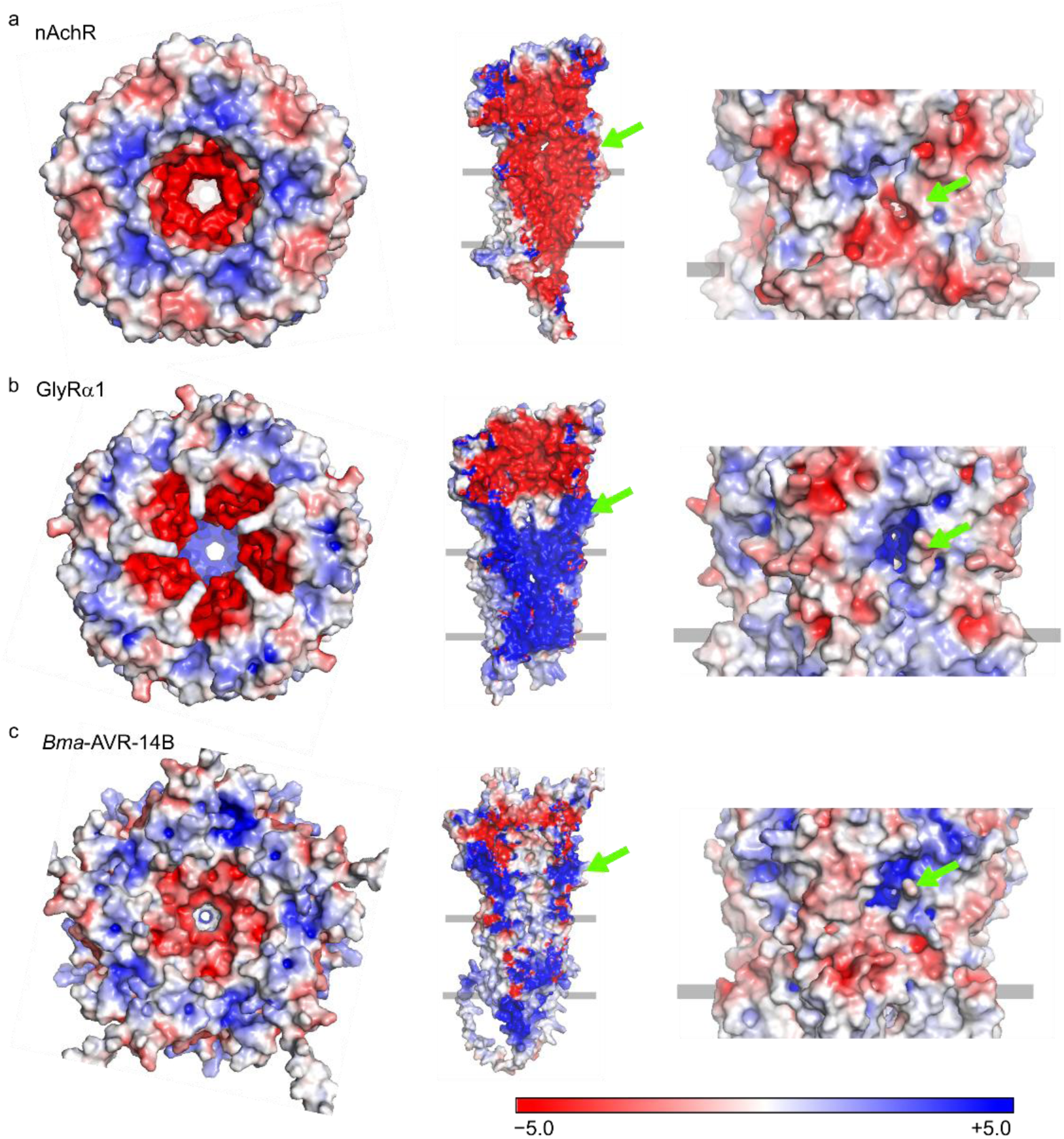
Electrostatic surface representations of Cys-loop receptors. Electrostatic potential maps are shown for the open-state human nAChR (PDB ID: 7KOX^53^) (**a**), the open-state zebrafish GlyRα1 (PDB ID: 6PM6^52^) (**b**), and the *Bma*-AVR-14B (**c**). For each receptor, three views are shown from left to right: a top view from the extracellular side; a side view of the pentamer highlighting the pore-facing surface with two subunits displayed for clarity; and a lateral view of the extracellular domain. Green arrows indicate a putative lateral pathway. Electrostatic potentials were calculated using APBS and mapped onto the molecular surface using a consistent color scale, where negative potentials (−5 k_B_T/e) are shown in red and positive potentials (+5 k_B_T/e) are shown in blue.

## References

1 Xiao, R. & Xu, X. Z. S. Temperature Sensation: From Molecular Thermosensors to Neural Circuits and Coding Principles. Annu Rev Physiol 83, 205–230 (2021). 10.1146/annurev-physiol-031220-095215

2 Dhaka, A., Viswanath, V. & Patapoutian, A. Trp ion channels and temperature sensation. Annu Rev Neurosci 29, 135–161 (2006). 10.1146/annurev.neuro.29.051605.112958

3 Castillo, K., Diaz-Franulic, I., Canan, J., Gonzalez-Nilo, F. & Latorre, R. Thermally activated TRP channels: molecular sensors for temperature detection. Phys Biol 15, 021001 (2018). 10.1088/1478-3975/aa9a6f

4 Ohnishi, K. et al. OSM-9 and OCR-2 TRPV channels are accessorial warm receptors in *Caenorhabditis elegans* temperature acclimatisation. Sci Rep 10, 18566 (2020). 10.1038/s41598-020-75302-3

5 Kumar Mondal, A., Carrillo, E., Jayaraman, V. & Twomey, E. C. Glutamate gating of AMPA-subtype iGluRs at physiological temperatures. Nature 641, 788–796 (2025). 10.1038/s41586-025-08770-0

6 Wolstenholme, A. J. Glutamate-gated chloride channels. J Biol Chem 287, 40232–40238 (2012). 10.1074/jbc.R112.406280

7 Vassilatis, D. K. et al. Evolutionary relationship of the ligand-gated ion channels and the avermectin-sensitive, glutamate-gated chloride channels. J Mol Evol 44, 501–508 (1997). 10.1007/pl00006174

8 Mizzi, N. & Blundell, R. Glycine receptors: Structure, function, and therapeutic implications. Molecular Aspects of Medicine 103, 101360 (2025). 10.1016/j.mam.2025.101360

9 Martin, R. J., Robertson, A. P. & Choudhary, S. Ivermectin: An Anthelmintic, an Insecticide, and Much More. Trends Parasitol 37, 48–64 (2021). 10.1016/j.pt.2020.10.005

10 Campbell, W. C., Fisher, M. H., Stapley, E. O., Albers-Schönberg, G. & Jacob, T. A. Ivermectin: A Potent New Antiparasitic Agent. Science 221, 823–828 (1983). doi:10.1126/science.6308762

11 Burg, R. W. et al. Avermectins, new family of potent anthelmintic agents: producing organism and fermentation. Antimicrob Agents Chemother 15, 361–367 (1979). 10.1128/aac.15.3.361

12 Cully, D. F. et al. Cloning of an avermectin-sensitive glutamate-gated chloride channel from *Caenorhabditis elegans*. Nature 371, 707–711 (1994). 10.1038/371707a0

13 Cully, D. F., Wilkinson, H., Vassilatis, D. K., Etter, A. & Arena, J. P. Molecular biology and electrophysiology of glutamate-gated chloride channels of invertebrates. Parasitology 113 Suppl, S191-200 (1996). 10.1017/s0031182000077970

14 Wolstenholme, A. J. & Rogers, A. T. Glutamate-gated chloride channels and the mode of action of the avermectin/milbemycin anthelmintics. Parasitology 131 Suppl, S85-95 (2005). 10.1017/s0031182005008218

15 Sultan, N. et al. Characterizing the Ion-Conductive State of the α7-Nicotinic Acetylcholine Receptor via Single-Channel Measurements and Molecular Dynamics Simulations. bioRxiv (2025). 10.1101/2025.08.15.670429

16 Cerdan, A. H., Peverini, L., Changeux, J. P., Corringer, P. J. & Cecchini, M. Lateral fenestrations in the extracellular domain of the glycine receptor contribute to the main chloride permeation pathway. Sci Adv 8, eadc9340 (2022). 10.1126/sciadv.adc9340

17 Miller, P. S. & Aricescu, A. R. Crystal structure of a human GABAA receptor. Nature 512, 270–275 (2014). 10.1038/nature13293

18 Laverty, D. et al. Cryo-EM structure of the human α1β3γ2 GABAA receptor in a lipid bilayer. Nature 565, 516–520 (2019). 10.1038/s41586-018-0833-4

19 Cook, A., et al. *Caenorhabditis elegans* ivermectin receptors regulate locomotor behaviour and are functional orthologues of *Haemonchus contortus* receptors. Molecular and Biochemical Parasitology 147, 118–125 (2006). 10.1016/j.molbiopara.2006.02.003

20 Lamassiaude, N., Courtot, E., Corset, A., Charvet, C. L. & Neveu, C. Pharmacological characterization of novel heteromeric GluCl subtypes from *Caenorhabditis elegans* and parasitic nematodes. Br J Pharmacol 179, 1264–1279 (2022). 10.1111/bph.15703

21 Moreno, Y., Nabhan, J. F., Solomon, J., Mackenzie, C. D. & Geary, T. G. Ivermectin disrupts the function of the excretory-secretory apparatus in microfilariae of *Brugia malayi*. Proc Natl Acad Sci U S A 107, 20120–20125 (2010). 10.1073/pnas.1011983107

22 Choudhary, S. et al. Nodulisporic acid produces direct activation and positive allosteric modulation of AVR-14B, a glutamate-gated chloride channel from adult *Brugia malayi*. Proceedings of the National Academy of Sciences 119, e2111932119 (2022). doi:10.1073/pnas.2111932119

23 Voets, T. et al. The principle of temperature-dependent gating in cold- and heat-sensitive TRP channels. Nature 430, 748–754 (2004). 10.1038/nature02732

24 Sakkas, H. & Papadopoulou, C. Antimicrobial Activity of Basil, Oregano, and Thyme Essential Oils. J Microbiol Biotechnol 27, 429–438 (2017). 10.4014/jmb.1608.08024

25 Gavaric, N., Mozina, S. S., Kladar, N. & Bozin, B. Chemical Profile, Antioxidant and Antibacterial Activity of Thyme and Oregano Essential Oils, Thymol and Carvacrol and Their Possible Synergism. Journal of Essential Oil Bearing Plants 18, 1013–1021 (2015). 10.1080/0972060X.2014.971069

26 Lynagh, T., Cromer, B. A., Dufour, V. & Laube, B. Comparative pharmacology of flatworm and roundworm glutamate-gated chloride channels: Implications for potential anthelmintics. Int J Parasitol Drugs Drug Resist 4, 244–255 (2014). 10.1016/j.ijpddr.2014.07.004

27 Grewer, C. Investigation of the α1-Glycine Receptor Channel-Opening Kinetics in the Submillisecond Time Domain. Biophysical Journal 77, 727–738 (1999). 10.1016/S0006-3495(99)76927-4

28 Inoue, K., Ueno, S., Yamada, J. & Fukuda, A. Characterization of newly cloned variant of rat glycine receptor α1 subunit. Biochemical and Biophysical Research Communications 327, 300–305 (2005). 10.1016/j.bbrc.2004.12.010

29 Zevian, S. C. & Yanowitz, J. L. Methodological considerations for heat shock of the nematode *Caenorhabditis elegans*. Methods 68, 450–457 (2014). 10.1016/j.ymeth.2014.04.015

30 Dent, J. A., Smith, M. M., Vassilatis, D. K. & Avery, L. The genetics of ivermectin resistance in *Caenorhabditis elegans*. Proceedings of the National Academy of Sciences 97, 2674–2679 (2000). doi:10.1073/pnas.97.6.2674

31 Shahi, K. & Baudry, M. Increasing binding affinity of agonists to glutamate receptors increases synaptic responses at glutamatergic synapses. Proc Natl Acad Sci U S A 89, 6881–6885 (1992). 10.1073/pnas.89.15.6881

32 Tocco, G., Massicotte, G., Standley, S., Thompson, R. F. & Baudry, M. Effect of Temperature and Calcium on the Binding Properties of the AMPA Receptor in Frozen Rat Brain Sections. Eur J Neurosci 4, 1093–1103 (1992). 10.1111/j.1460-9568.1992.tb00136.x

33 Schmiedhofer, P., Vogel, F. D., Koniuszewski, F. & Ernst, M. Cys-loop receptors on cannabinoids: All high? Front Physiol 13, 1044575 (2022). 10.3389/fphys.2022.1044575

34 Olsen, R. W. et al. Structural models of ligand-gated ion channels: sites of action for anesthetics and ethanol. Alcohol Clin Exp Res 38, 595–603 (2014). 10.1111/acer.12283

35 Jalalypour, F., Howard, R. J. & Lindahl, E. Allosteric Cholesterol Site in Glycine Receptors Characterized through Molecular Simulations. The Journal of Physical Chemistry B 128, 4996–5007 (2024). 10.1021/acs.jpcb.4c01703

36 O’Lone, R. B. & Campbell, W. C. Effect of refrigeration on the antinematodal efficacy of ivermectin. J Parasitol 87, 452–454 (2001). 10.1645/0022-3395(2001)087[0452:eorota]2.0.co;2

37 Kavallieratos, N. G., Athanassiou, C. G., Vayias, B. J., Mihail, S. B. & Tomanović, Z. Insecticidal efficacy of abamectin against three stored-product insect pests: influence of dose rate, temperature, commodity, and exposure interval. J Econ Entomol 102, 1352–1359 (2009). 10.1603/029.102.0363

38 Esquivel-Román, A., Baena-Díaz, F., Bustos-Segura, C., De Gasperin, O. & González-Tokman, D. Synergistic effects of elevated temperature with pesticides on reproduction, development and survival of dung beetles. Ecotoxicology 34, 207–218 (2025). 10.1007/s10646-024-02825-0

39 Nie, Y. et al. Combined effects of abamectin and temperature on the physiology and behavior of male lizards (Eremias argus): Clarifying adaptation and maladaptation. Sci Total Environ 837, 155794 (2022). 10.1016/j.scitotenv.2022.155794

40 Lin, C. et al. Molecular and circuit mechanisms underlying avoidance of rapid cooling stimuli in *C. elegans*. Nature Communications 15, 297 (2024). 10.1038/s41467-023-44638-5

41 Gat, A. et al. Integration of spatially opposing cues by a single interneuron guides decision-making in *C. elegans*. Cell Rep 42, 113075 (2023). 10.1016/j.celrep.2023.113075

42 Li, Z. et al. A *C. elegans* neuron both promotes and suppresses motor behavior to fine tune motor output. Front Mol Neurosci 16, 1228980 (2023). 10.3389/fnmol.2023.1228980

43 Hammarlund, M., Hobert, O., Miller, D. M., 3rd & Sestan, N. The CeNGEN Project: The Complete Gene Expression Map of an Entire Nervous System. Neuron 99, 430–433 (2018). 10.1016/j.neuron.2018.07.042

44 Ohnishi, K. et al. G protein-coupled receptor-based thermosensation determines temperature acclimatization of *Caenorhabditis elegans*. Nature Communications 15, 1660 (2024). 10.1038/s41467-024-46042-z

45 Glauser, D. A. Temperature sensing and context-dependent thermal behavior in nematodes. Current Opinion in Neurobiology 73, 102525 (2022). 10.1016/j.conb.2022.102525

46 Rubio-Tomás, T., Alegre-Cortés, E., Lionaki, E., Fuentes, J. M. & Tavernarakis, N. Heat shock and thermotolerance in *Caenorhabditis elegans*: An overview of laboratory techniques. Methods Cell Biol 185, 1–17 (2024). 10.1016/bs.mcb.2024.02.001

47 Ghit, A., Assal, D., Al-Shami, A. S. & Hussein, D. E. E. GABA(A) receptors: structure, function, pharmacology, and related disorders. J Genet Eng Biotechnol 19, 123 (2021). 10.1186/s43141-021-00224-0

48 San Martín, V. P., Sazo, A., Utreras, E., Moraga-Cid, G. & Yévenes, G. E. Glycine Receptor Subtypes and Their Roles in Nociception and Chronic Pain. Front Mol Neurosci 15, 848642 (2022). 10.3389/fnmol.2022.848642

## References

49 Fujiwara, Y., Keceli, B., Nakajo, K. & Kubo, Y. Voltage- and [ATP]-dependent Gating of the P2X2 ATP Receptor Channel. Journal of General Physiology 133, 93–109 (2008). 10.1085/jgp.200810002

50 Fujiwara, Y. et al. The cytoplasmic coiled-coil mediates cooperative gating temperature sensitivity in the voltage-gated H+ channel Hv1. Nature Communications 3, 816 (2012). 10.1038/ncomms1823

51 Abramson, J. et al. Accurate structure prediction of biomolecular interactions with AlphaFold 3. Nature 630, 493–500 (2024). 10.1038/s41586-024-07487-w

52 Yu, J. et al. Mechanism of gating and partial agonist action in the glycine receptor. Cell 184, 957–968.e921 (2021). 10.1016/j.cell.2021.01.026

53 Noviello, C. M. et al. Structure and gating mechanism of the α7 nicotinic acetylcholine receptor. Cell 184, 2121–2134.e2113 (2021). 10.1016/j.cell.2021.02.049

54 Brenner, S. The genetics of *Caenorhabditis elegans*. Genetics 77, 71–94 (1974). 10.1093/genetics/77.1.71

55 Ohta, A. et al. The intron binding protein EMB-4 is an opposite regulator of cold and high temperature tolerance in *Caenorhabditis elegans*. PNAS Nexus 3 (2024). 10.1093/pnasnexus/pgae293

56 Vriens, J., Nilius, B. & Voets, T. Peripheral thermosensation in mammals. Nature Reviews Neuroscience 15, 573–589 (2014). 10.1038/nrn3784

57 Elias, M., Wieczorek, G., Rosenne, S. & Tawfik, D. S. The universality of enzymatic rate–temperature dependency. Trends in Biochemical Sciences 39, 1–7 (2014). 10.1016/j.tibs.2013.11.001

58 Hille, B., "Ionic Channels of Excitable Membranes", 3rd ed. (Sunderland, MA: Sinauer Associates, Inc., 2001).

59 Kanda, Y. Investigation of the freely available easy-to-use software ‘EZR’ for medical statistics. Bone Marrow Transplantation 48, 452–458 (2013). 10.1038/bmt.2012.244

